# A Comprehensive Assessment of Human Natural Killer Cell Phenotype and Function in Whole Blood

**DOI:** 10.1101/2020.02.03.932640

**Authors:** M Market, G Tennakoon, J Ng, M Scaffidi, M A Kennedy, C Tanese de Souza, R C Auer

**Affiliations:** University of Ottawa, Ottawa, ON, Canada; The Ottawa Hospital Research Institute, Ottawa, ON, Canada; Department of Surgery, University of Ottawa, The Ottawa Hospital, Ottawa, ON, Canada

**Keywords:** Natural Killer cells, whole blood, flow cytometry, cytokine stimulation, interferon gamma, NKG2D

## Abstract

The majority of data on human Natural Killer (NK) cell phenotype and function has been generated using cryopreserved peripheral blood mononuclear cells (PBMCs). However, cryopreservation can have adverse effects on PBMCs. In contrast, investigating immune cells in whole blood can reduce the time, volume of blood required, and potential artifacts associated with manipulation of the cells. Whole blood collected from healthy donors and cancer patients was processed by three separate protocols that can be used independently or in parallel to assess extracellular receptors, intracellular signaling protein phosphorylation, and intracellular and extracellular cytokine production in human NK cells. To assess extracellular receptor expression, 200 μL of whole blood was incubated with an extracellular staining (ECS) mix and cells were subsequently fixed and RBCs lysed prior to analysis. The phosphorylation status of signaling proteins was assessed in 500 μL of whole blood following co-incubation with interleukin (IL)−2/12 and the ECS mix for 20 minutes prior to cell fixation and RBC lysis and subsequent permeabilization for staining with an intracellular staining (ICS) mix. Cytokine production (IFNγ) was similarly assessed by incubating 1 mL of whole blood with PMA-ionomycin or IL−2/12 prior to incubation with ECS and subsequent ICS antibodies. In addition, plasma was collected from stimulated samples prior to ECS for quantification of secreted IFNγ by ELISA. Results were consistent, despite inherent interpatient variability. Although we did not investigate an exhaustive list of targets, this approach enabled quantification of representative ECS surface markers including activating (NKG2D and DNAM−1) and inhibitory (NKG2D, PD−1, TIGIT, and TIM−3) receptors, cytokine receptors (CD25, CD122, CD132, and CD212) and ICS markers associated with NK cell activation following stimulation, including signaling protein phosphorylation (p-STAT4, p-STAT5, p-p38 MAPK, p-S6K) and IFNγ in both healthy donors and cancer patients. In addition, we compared extracellular receptor expression using whole blood versus cryopreserved PBMCs and observed a significant difference in the expression of almost all receptors. The methods presented permit a relatively rapid parallel assessment of immune cell receptor expression, signaling protein activity, and cytokine production in a minimal volume of whole blood from both healthy donors and cancer patients.

## 1 Introduction

Natural Killer (NK) cells, first identified by Kiessling et al. in 1975, are cytotoxic lymphocytes that play a critical role in the innate immune response through the destruction of stressed, infected, or cancerous cells(1). Defective NK cell function has been linked to autoimmune and infectious diseases as well as cancer(2–6). Our investigations focus on understanding the suppression of NK cells following surgery in cancer patients and the impact of immunosuppression on metastasis. Specifically, our lab and others have shown that postoperative defects in NK cell cytotoxicity and IFNγ production contribute to increased metastasis in models of surgical stress(7–9). Our initial observations of this suppressed phenotype were in cryopreserved PBMCs; however, we have also observed this phenomenon in whole blood. We then developed protocols that can be used in parallel to assess the phenotype, intracellular signaling following cytokine stimulation, and cytokine production of immune cells, and as an example, in this paper we highlight its implementation for our ongoing research investigating NK cells in cancer patients.

For practical reasons, the majority of the data on human NK cells has been generated using peripheral blood mononuclear cells (PBMCs). For instance, cryopreservation allows for running batched samples simultaneously as well as logistical flexibility for the storage and shipment of samples between research facilities(10). Using this approach, the study of cryopreserved PBMCs through functional and phenotypic assays has yielded a great deal of understanding about the role of NK cell function in disease. However, the use of cryopreserved PBMCs in immunologic studies is associated with adverse effects on cell populations/ certain cell markers and altered gene expression(11–13). As a result, our understanding of NK cells may benefit in certain circumstances from investigations of non-cryopreserved cells.

In trying to assess the mechanism of NK cell dysfunction in cancer patients in the context of surgery, we sought to assess key markers and intracellular pathways associated with this dysfunctional NK cell phenotype. We investigated upstream receptor expression and subsequent signaling protein phosphorylation in order to elucidate the mechanism of NK cell suppression. NK cells do not undergo clonal selection, they instead express a limited number of germline-encoded receptors(14). NK cell activating receptors recognize pathogen-derived antigens as well as stress-induced ligands in what is termed the “induced-self recognition model”(15–17). These activating signals are antagonized by inhibitory receptors that recognize constitutively expressed self-molecules or inhibitory checkpoint proteins(15,16). We sought to assess the expression levels of the activating receptors NKG2D and DNAM−1 and the inhibitory receptors NKG2A, PD−1, TIGIT, and TIM−3. In addition to these receptors, NK cells also express a plethora of cytokine receptors, including interleukin (IL)−2R and IL−12R(18). NK cell activity is thus regulated by the integration of activating and inhibitory ligands through these many receptors, which results in phosphorylation and transduction through signaling proteins such as STAT4, STAT5, p38 MAPK, and S6K(9,19–23). This culminates in the regulation of transcription factor activity that controls the transcription of cytokines such as IFNγ and cytotoxic proteins, including granzymes and perforin(24,25). In characterizing the perioperative NK cell phenotype, we found it challenging to assess phosphorylation status in cryopreserved PBMCs. As a solution, we considered the use of whole blood, which proved to be far superior. In the troubleshooting process we also discovered a discrepancy between the phenotypes observed in cryopreserved PBMCs versus whole blood staining. The successes we experienced by using whole blood samples, compared to cryopreserved PBMCs, prompted us to continue using whole blood samples for assessment of NK cell activity and develop a series of easily implemented, standardized protocols that enable a comprehensive investigation of NK phenotype and function.

There is a paucity of studies investigating immune cell function from whole blood(26). We posit that such studies would avoid the adverse effects of cryopreservation and provide more biologically relevant results in some circumstances. For example, investigating protein phosphorylation states by flow cytometry is difficult in cryopreserved samples due to the low signal to noise ratio of the target protein compared to investigations in whole blood samples(27). Many of these limitations can be overcome by staining directly in whole blood, which also allows for simpler and faster protocols that require minimal manipulation of the cells of interest and therefore support the biological relevance of the results. A limitation of whole blood assays includes having to process patient samples immediately and therefore they cannot be tested simultaneously, which could lead to greater inter-assay variability. However, technical expertise, appropriate controls, and validated standard operating procedures can be implemented to help mitigate this limitation.

Comparisons of immunologic assays using cryopreserved PBMCs and whole blood samples have previously been reported and is not the focus of our report(24,25,28). Here we sought to highlight the feasibility and advantages of using whole blood samples as a strategy for phenotypic and functional assessments in NK cells. As a proof of concept, we show the utilization of these protocols in our ongoing research. We explored the differential expression of phenotypic receptors necessary for NK activity and phosphorylation of downstream signaling molecules in healthy donors and cancer patients using whole blood. Finally, NK cell function was investigated by quantifying intracellular and extracellular IFNγ by flow cytometry and ELISA following stimulation with IL−2/IL−12 or PMA/Ionomycin. We show that assaying cryopreserved cells results in altered NK cell phenotype in human patients as compared to whole blood analysis. In addition, we outline in detail novel whole blood protocols that can be used in parallel to assess immune cell receptor expression, signaling protein phosphorylation, and cytokine production. Although developed to assess NK cell activity in the perioperative period, these protocols could be used to assess other immune cell phenotypes in other pathological conditions.

## 2 Materials and Equipment

- 37°C incubator
- 37°C water bath
- Centrifuge
- Sodium-heparin tubes (BD Vacutainer® Cat #367878/ 367874)
- BD FACS Lyse/ Fix Buffer (Cat #558049)
- Deionized/ distilled H_2_O
- BD Perm III Buffer (Cat #558050)
- 1% Paraformaldehyde
- Recombinant human IL−2 (Tecin Teceleukin)
- Recombinant human IL−12 (R&D System Cat #219-IL005)
- Phosphate Buffered Saline (PBS)
- Flow Buffer (PBS + 2.5g BSA + 0.5M EDTA)
- Biolegend Human Fc block (Cat #422302)
- BD Golgi plug (Brefaldin A) (Cat #51−2301K2)
- R&D Quantikine Human IFNγ ELISA (Cat #DIF50)
- Flow cytometer (LSR Fortessa)
- Antibodies: (**Table 1**)

**Table 1.**
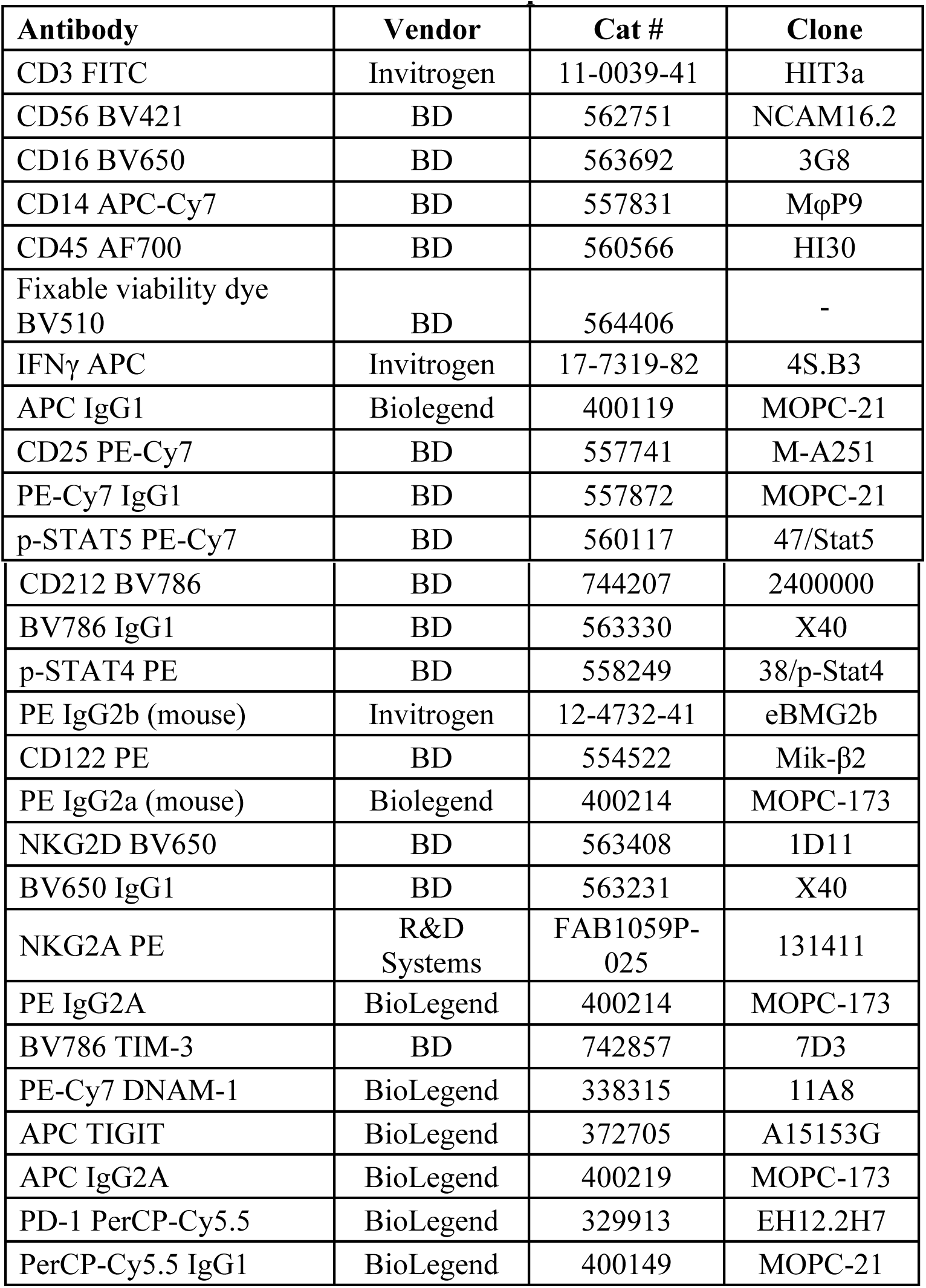
Antibodies used in whole blood panels.

| **Antibody** | **Vendor** | **Cat #** | **Clone** |
| --- | --- | --- | --- |
| CD3 FITC | Invitrogen | 11-0039-41 | HIT3a |
| CD56 BV421 | BD | 562751 | NCAM16.2 |
| CD16 BV650 | BD | 563692 | 3G8 |
| CD14 APC-Cy7 | BD | 557831 | MφP9 |
| CD45 AF700 | BD | 560566 | HI30 |
| Fixable viability dye BV510 | BD | 564406 | - |
| IFNγ APC | Invitrogen | 17-7319-82 | 4S.B3 |
| APC IgG1 | Biolegend | 400119 | MOPC-21 |
| CD25 PE-Cy7 | BD | 557741 | M-A251 |
| PE-Cy7 IgG1 | BD | 557872 | MOPC-21 |
| p-STAT5 PE-Cy7 | BD | 560117 | 47/Stat5 |
| CD212 BV786 | BD | 744207 | 2400000 |
| BV786 IgG1 | BD | 563330 | X40 |
| p-STAT4 PE | BD | 558249 | 38/p-Stat4 |
| PE IgG2b (mouse) | Invitrogen | 12-4732-41 | eBMG2b |
| CD122 PE | BD | 554522 | Mik-β2 |
| PE IgG2a (mouse) | Biolegend | 400214 | MOPC-173 |
| NKG2D BV650 | BD | 563408 | 1D11 |
| BV650 IgG1 | BD | 563231 | X40 |
| NKG2A PE | R&D Systems | FAB1059P-025 | 131411 |
| PE IgG2A | BioLegend | 400214 | MOPC-173 |
| BV786 TIM-3 | BD | 742857 | 7D3 |
| PE-Cy7 DNAM-1 | BioLegend | 338315 | 11A8 |
| APC TIGIT | BioLegend | 372705 | A15153G |
| APC IgG2A | BioLegend | 400219 | MOPC-173 |
| PD-1 PerCP-Cy5.5 | BioLegend | 329913 | EH12.2H7 |
| PerCP-Cy5.5 IgG1 | BioLegend | 400149 | MOPC-21 |

## 3 Methods

This protocol was approved by the Ottawa Health Science Research Ethics Board. All subjects gave written informed consent in accordance with the Declaration of Helsinki. Eligible patients were >18 years of age and had a planned surgical resection of the primary or metastatic tumour (cancer patients) or healthy donors who volunteered to participate. Exclusion criteria included a history of active viral or bacterial infection or known HIV or Hepatitis B or C, autoimmune diseases, or use of immunosuppressive medications.

### 3.1 Protocol 1 – Extracellular Receptor Staining (Figure 1A)

Objective: Assess cell surface receptor expression in Natural Killer cells from whole blood.

1. Collect whole blood in sodium-heparin tubes (blood drawn by trained nurse/ phlebotomist)
2. Prepare FACS lyse/ fix buffer in advance (1:5 dilution with diH_2_O; 20X blood volume per sample)
3. Transfer 200 μL of whole blood to 15 mL falcon tube
4. Add extracellular staining (ECS) mix (40 μL); Mix by pipetting up and down
5. Incubate for 15 mins at room temperature
6. Add FACS lyse/ fix buffer
7. Incubate for 10 minutes in 37°C water bath
8. Spin at 500 g for 8 minutes
9. Aspirate supernatant and resuspend in 1 mL flow buffer
10. Spin at 500 g for 5 mins
11. Resuspend in 200 μL flow buffer
12. Store at 4°C for up to 24 hours

### 3.2 Protocol 2 – Intracellular Signaling Protein Phosphorylation Staining (Figure 4A)

Objective: Assess signaling protein/ transcription factor phosphorylation in response to stimuli in Natural Killer cells from whole blood.

1. Collect whole blood in sodium-heparin tubes (blood drawn by trained nurse/ phlebotomist)
2. Prepare FACS lyse/ fix buffer in advance (1:5 dilution with diH_2_O; 20X blood volume per sample)
3. Aliquot 500 μL blood into new sodium-heparin tubes
4. Add stimulation and ECS mix and incubate for 20 minutes in 37°C water bath
5. Transfer whole blood to 15 mL falcon tubes
6. Add FACS lyse/fix buffer
7. Incubate for 10 minutes in 37°C water bath
8. Spin at 500 g for 8 minutes
9. Aspirate supernatant and wash with 1 mL flow buffer
10. Spin at 500 g for 5 minutes
11. Aspirate supernatant and resuspend in 500 μL chilled BD Perm III buffer
12. Incubate on ice in the dark for 30 minutes
13. Spin at 300 g for 10 minutes
14. Aspirate supernatant and resuspend in 400 μL flow buffer
15. Transfer 200 μL/well into 96 well v-bottom plate
16. Spin at 500 g for 5 minutes
17. Empty plate and resuspend in appropriate intracellular staining (ICS) mix (200 μL)
18. Incubate at room temperature in the dark for 1 hour
19. Spin at 500 g for 5 mins
20. Empty plate and resuspend in 200 μL flow buffer
21. Store at 4°C for up to 24 hours

### 3.3 Protocol 3 – Intracellular IFNγ Staining (Figure 5A)

Objective: Quantify intracellular IFNγ production as a measure of activity in Natural Killer cells from whole blood.

1. Collect whole blood in sodium-heparin tubes (blood drawn by trained nurse/ phlebotomist)
2. Prepare FACS lyse/ fix buffer in advance (1:5 dilution with diH_2_O; 20X blood volume per sample)
3. Aliquot 1 mL of whole blood into new sodium-heparin tubes
4. Incubate whole blood with PMA-ionomycin for 5 hours at 37°C and IL−2/IL−12 for 24 hours at 37°C
5. Add 10 ug/mL Golgi plug (Brefaldin A) per tube for the last 2 hours of each incubation
6. Invert tubes 10 times and incubate at 37°C for remaining 2 hours
7. Collect 600 μL whole blood in Eppendorf tube, spin at 13000 rpm for 1 minute, and store at −80°C for IFNγ ELISA
8. Transfer remaining 400 μL to 15mL falcon tubes
9. Incubate with Fc block (50 μL) for 5 mins at room temperature
10. Add ECS mix (40 μL); Mix by pipetting up and down
11. Incubate for 15 mins at room temperature
12. Add FACS lyse/ fix buffer
13. Incubate for 10 minutes in 37°C water bath
14. Spin at 500 g for 8 minutes
15. Aspirate supernatant and resuspend in 1 mL flow buffer
16. Spin at 500 g for 5 mins
17. Aspirate supernatant and resuspend in 500 μL chilled BD Perm III buffer
18. Incubate on ice in the dark for 30 minutes
19. Spin at 300 g for 10 minutes
20. Aspirate supernatant and resuspend in 400 μL flow buffer
21. Transfer 200 μL/ well into 96 well v-bottom plate
22. Spin at 500 g for 5 minutes
23. Empty plate and resuspend in appropriate ICS mix (200 μL)
24. Incubate at 4°C for 30 minutes
25. Empty plate and resuspend in 200 μL flow buffer
26. Store at 4°C for up to 24 hours

### Extracellular IFNγ Quantification (Figure 6A)

Objective: Quantify extracellular IFNγ production as a measure of activity in plasma from whole blood.

1. Thaw plasma samples at room temperature
2. Prior to beginning the ELISA, dilute samples at 1:5 or 1:0 for optimal quantification
3. Follow R&D Quantikine Human IFNγ ELISA protocol for patient plasma

## 4 Data Analysis

Descriptive statistics were used to summarize data collected on extracellular receptors, phospho-signaling proteins, and IFNγ production (median with interquartile range (IQR)). Wilcoxon matched-pairs signed rank test was used to determine if there were significant changes in receptor expression (percentage and MFI) between cryopreserved and whole blood samples. The level for statistical significance was set a priori at ≤0.05 (*p≤0.05, **p≤0.005, ***p≤0.0005, ****p≤0.00005). All statistical analyses were performed using Prism 8.

## 5 Results

### 5.1 Protocol 1 – Extracellular Receptor Staining

#### Quantifying NK cell surface receptors in whole blood

Using a 10 colour flow cytometry panel, we assessed the surface expression of six NK cell receptors, which are known to activate (NKG2D and DNAM−1) or inhibit (NKG2A, PD−1, TIGIT, and TIM−3) NK cell effector functions in a cohort of 16 healthy donors and 20 cancer patients (**Table 2**). Using a 9 colour flow cytometry panel, we similarly assessed whether the expression of IL−2/12 receptor subunits (CD25 (α), CD122 (β), CD132 (γ), and CD212 (β1)) could be detected in 13 healthy donors and 11 cancer patients (**Table 2**). We assessed the percentage of positive cells as well as the relative expression level (median fluorescence intensity/ MFI) of both activating/ inhibitory and cytokine receptors in NK cells (CD56^+^CD3^−^) using the indicated gating strategy (**Figure 1B and C**). We were able to assess both activating/ inhibitory and cytokine receptor expression using this whole blood protocol (**Figure 2A,B and Supplemental Table 1**).

**Figure 1.**
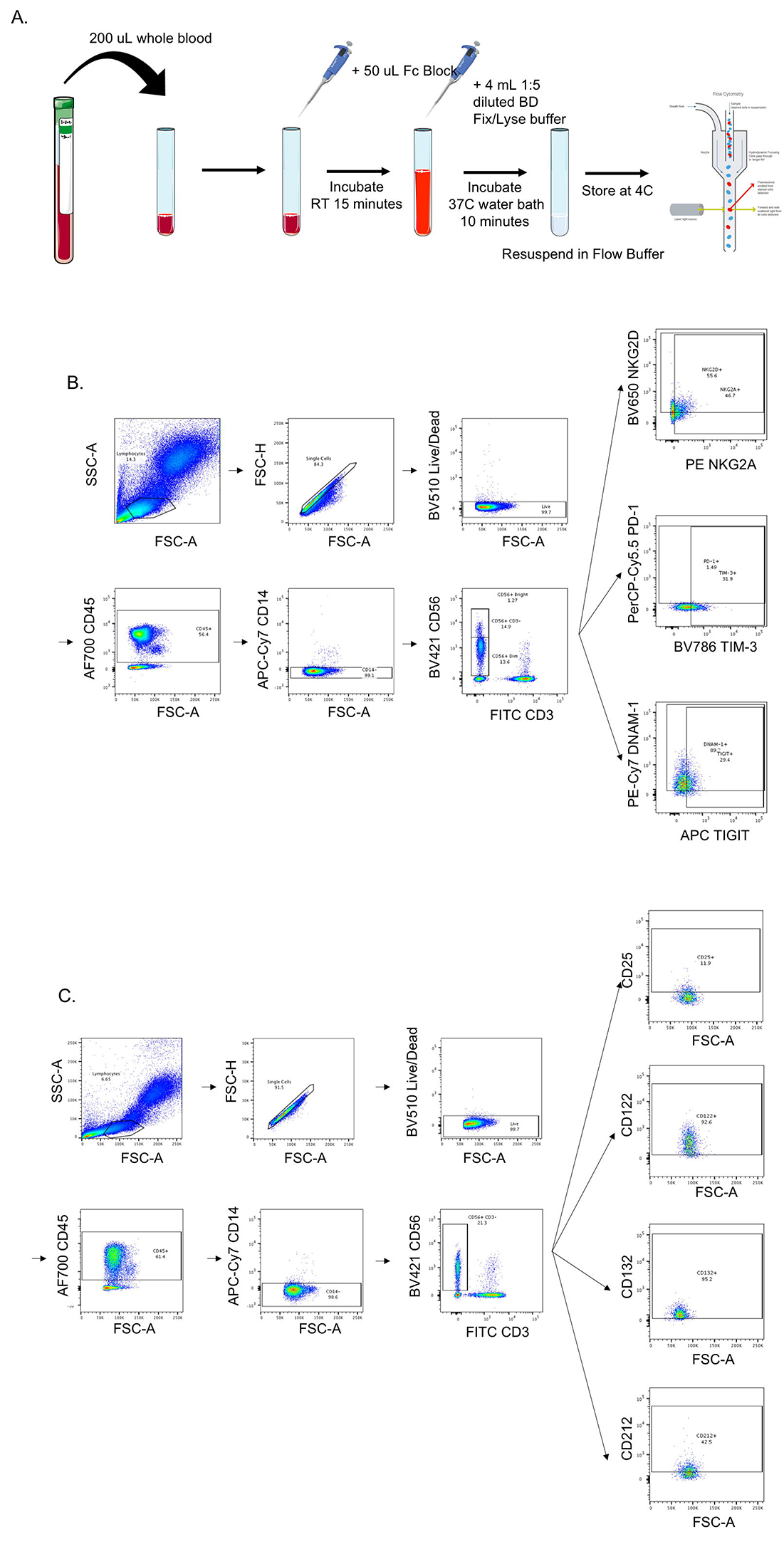
Whole blood receptor methodology and gating strategy. Whole blood was collected from patients and 200 μL was aliquoted per receptor panel. After a 15-minute incubation with the receptor panel, lyse/fix buffer was added and incubated for 10 minutes before blood was spun down, cells were washed and resuspended to be assessed by flow cytometry (**A**). The lymphocyte population was gated on before excluding doublets and dead cells. CD45^+^CD14^−^CD56^+^CD3^−^cells were gated on to assess activating/ inhibitory receptor (**B**) and cytokine receptor (**C**) expression based on isotype staining.

**Figure 2.**
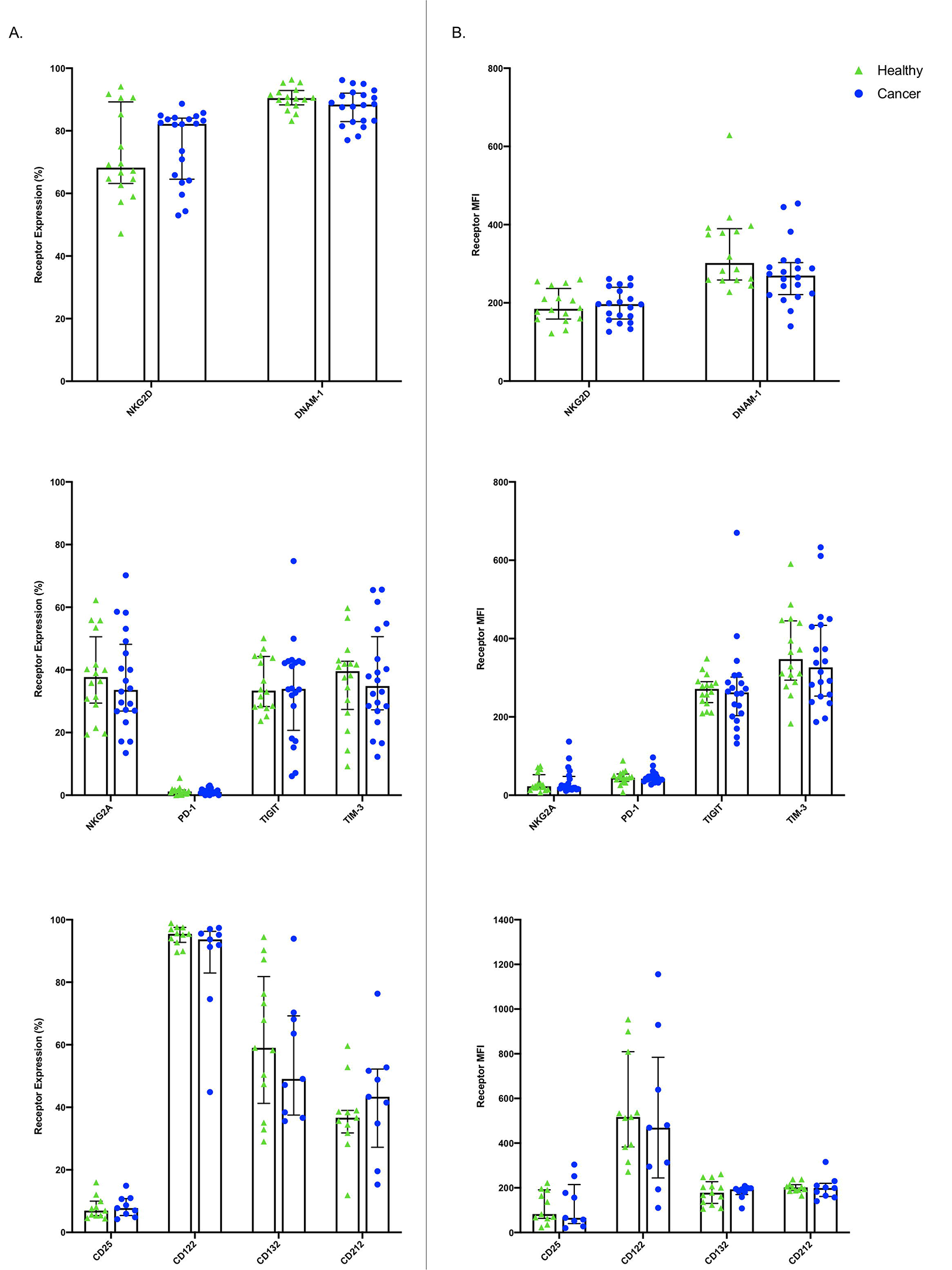
Whole blood extracellular surface receptor expression. The percentage of CD56+ CD3-cells expressing activating receptors (NKG2D and DNAM−1), inhibitory receptors (NKG2A, PD−1, TIGIT, and TIM−3), and cytokine receptor subunits (CD25, CD122, CD132, and CD212) was assessed in healthy donors (n=29) and cancer patients (n=31) (**A**). The relative level of expression (MFI) of the same activating/ inhibitory/ cytokine receptors was also assessed in CD56^+^CD3^−^cells (**B**). Shown are the median values ± IQR.

**Table 2.**
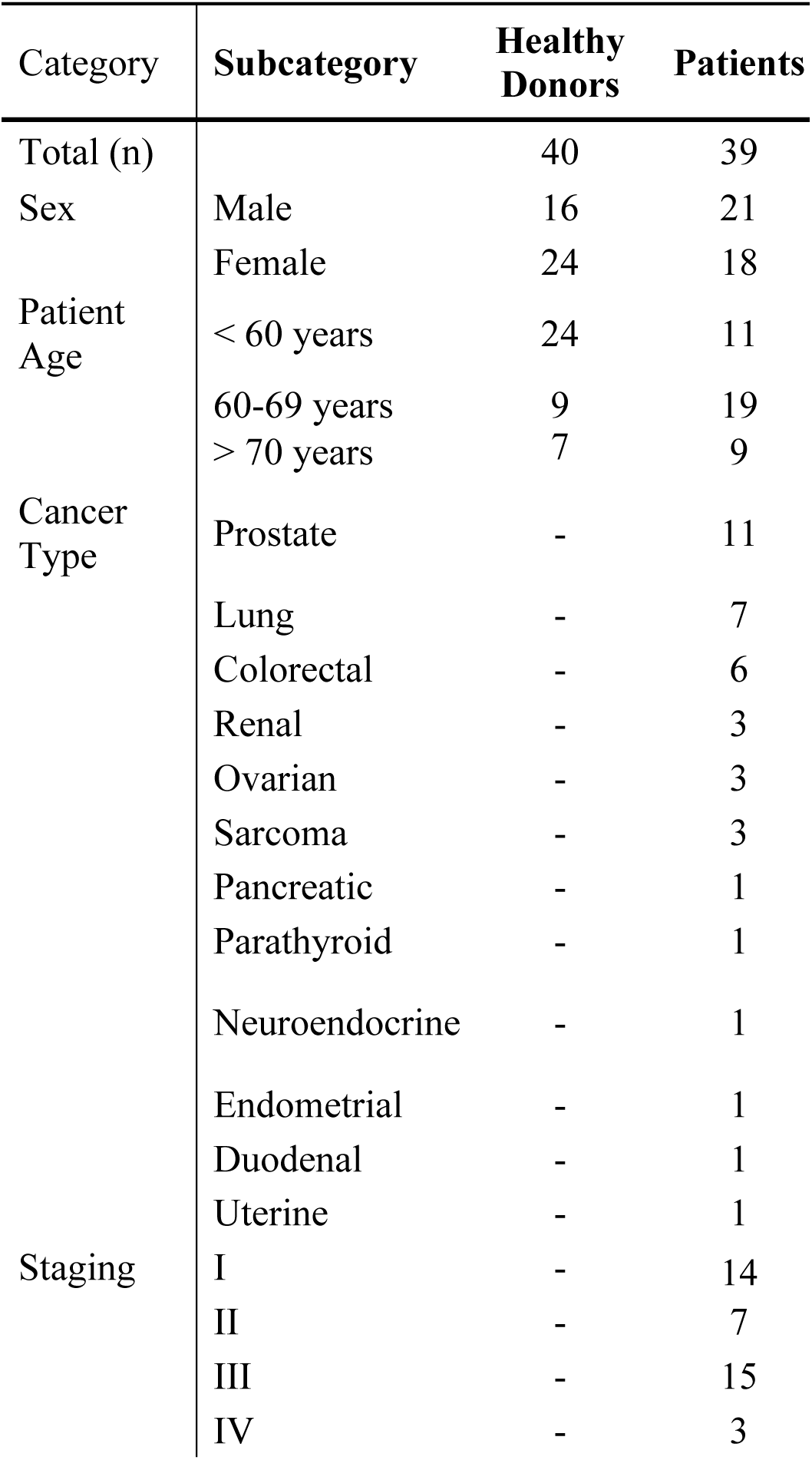
Whole blood patient demographics. Summary of patient information including sex, age, cancer type, and cancer staging for patient samples used to quantify receptor expression, phosphorylation status, and IFNγ production using whole blood.

**Table 3.** Cryopreserved PBMC patient demographics. Summary of patient information including sex, age, cancer type, and cancer staging for patient samples used to quantify activating/ inhibitory/ cytokine receptors using cryopreserved PBMCs.

| Category | Subcategory | Patients |
| --- | --- | --- |
| Total (n) |  | 22 |
| Gender | Male | 19 |
|  | Female | 6 |
| Patient Age | < 60 years | 7 |
|  | 60-69 years | 13 |
|  | > 70 years | 5 |
| Cancer Type | Prostate | 13 |
|  | Lung | 6 |
|  | Colorectal | 2 |
|  | Esophageal | 1 |
| Staging | I | 4 |
|  | II | 4 |
|  | III | 10 |
|  | IV | 1 |

#### Discrepancies between whole blood and cryopreserved NK cell surface receptors

In addition to whole blood we also assessed the expression of NKG2D, DNAM−1, PD−1, TIGIT, and TIM−3 (n=10) and CD25 and CD212 (n=11) in NK cells from cryopreserved PBMCs. After ficoll density centrifugation, PBMCs were isolated, washed, and stored in liquid nitrogen in 90% FBS 10% DMSO. We followed a standard protocol whereby PBMCs were thawed, rested overnight, and stained using a 10 colour (activating/ inhibitory receptors) or a 9 colour (cytokine receptors) flow cytometry panel(29–32). Consistent with previous publications, we found significant differences between the percentage of positive cells and receptor MFI in cryopreserved versus whole blood NK cells(33,34). (**Figure 3, Supplemental Table 2**).

**Figure 3.**
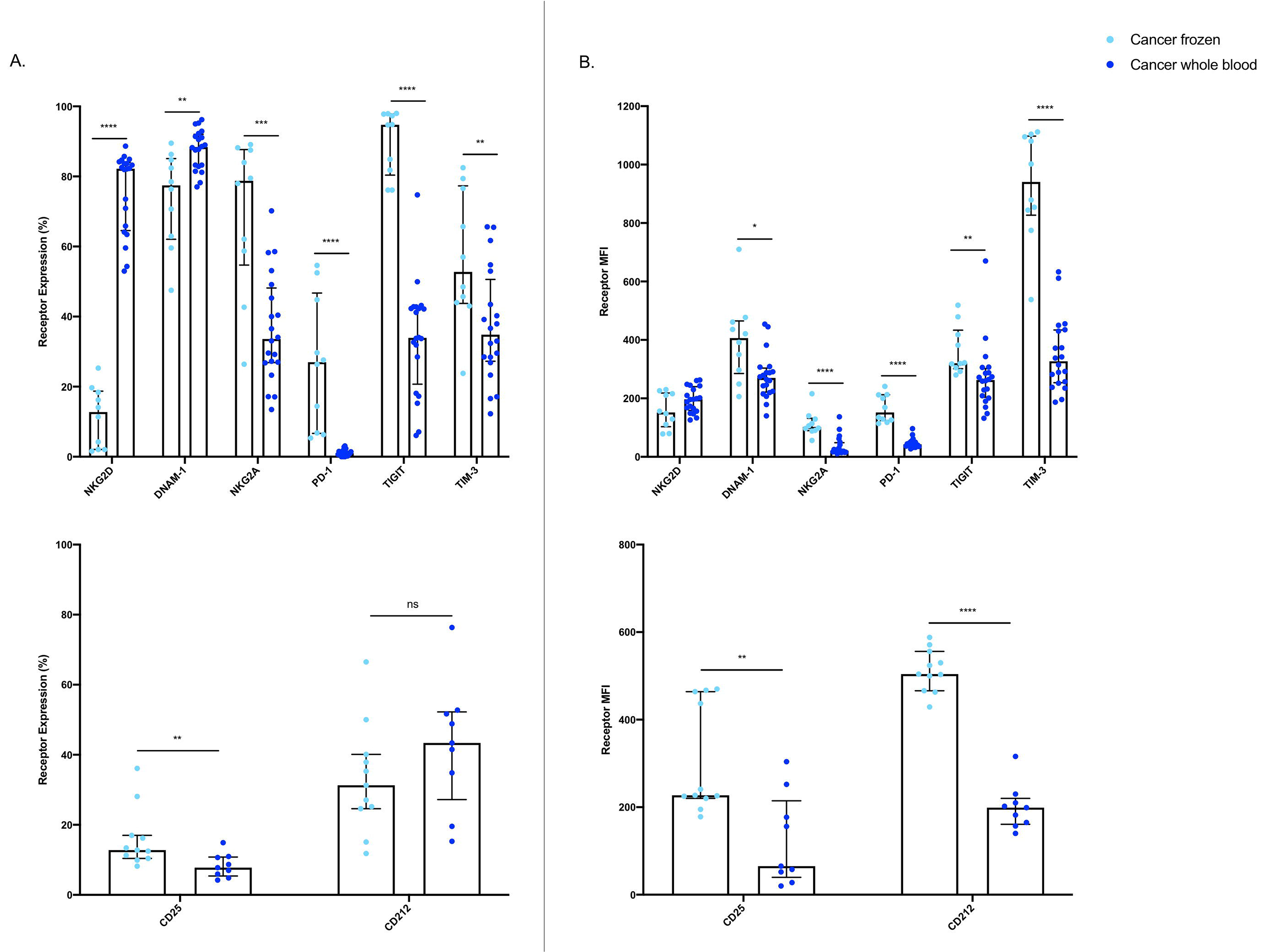
Expression of extracellular surface receptors differs between cryopreserved PBMCs and whole blood samples. A significant difference was observed between the percentage of CD56^+^CD3^−^cells expressing all activating and inhibitory receptors (cryopreserved n=10, whole blood n=20), as well as the cytokine receptor subunit CD25 (cryopreserved n=11, whole blood n=11) (**A**). In addition, a significant difference was also observed in the relative expression levels of DNAM−1, all inhibitory receptors, and both cytokine receptor subunits (**B**). Shown are the median values ± IQR. The Mann-Whitney test was used to assess statistical significance. p ≤ 0.05 (*p≤0.05, **p≤0.005, ***p≤0.0005, ****p≤0.00005).

### 5.2 Protocol 2 – Intracellular Signaling Protein Phosphorylation Staining

#### Detecting NK cell cytokine signaling in whole blood

The phosphorylation of signaling proteins downstream of IL−2/12 receptors was assessed in 13 healthy donors and 9 cancer patients. Activity was assessed with two 7 colour flow cytometry panels that included phospho-specific antibodies against STAT5, STAT4, p38 MAPK, and S6K. The indicated gating strategy allowed for the assessment of the relative expression level (MFI) of phosphorylated protein in NK cells (**Figure 4B**). We were able to quantify the phosphorylation of these signaling molecules in response to IL−2/12 stimulation in both healthy donors and cancer patients (**Figure 4C, Supplemental Table 3**).

**Figure 4.**
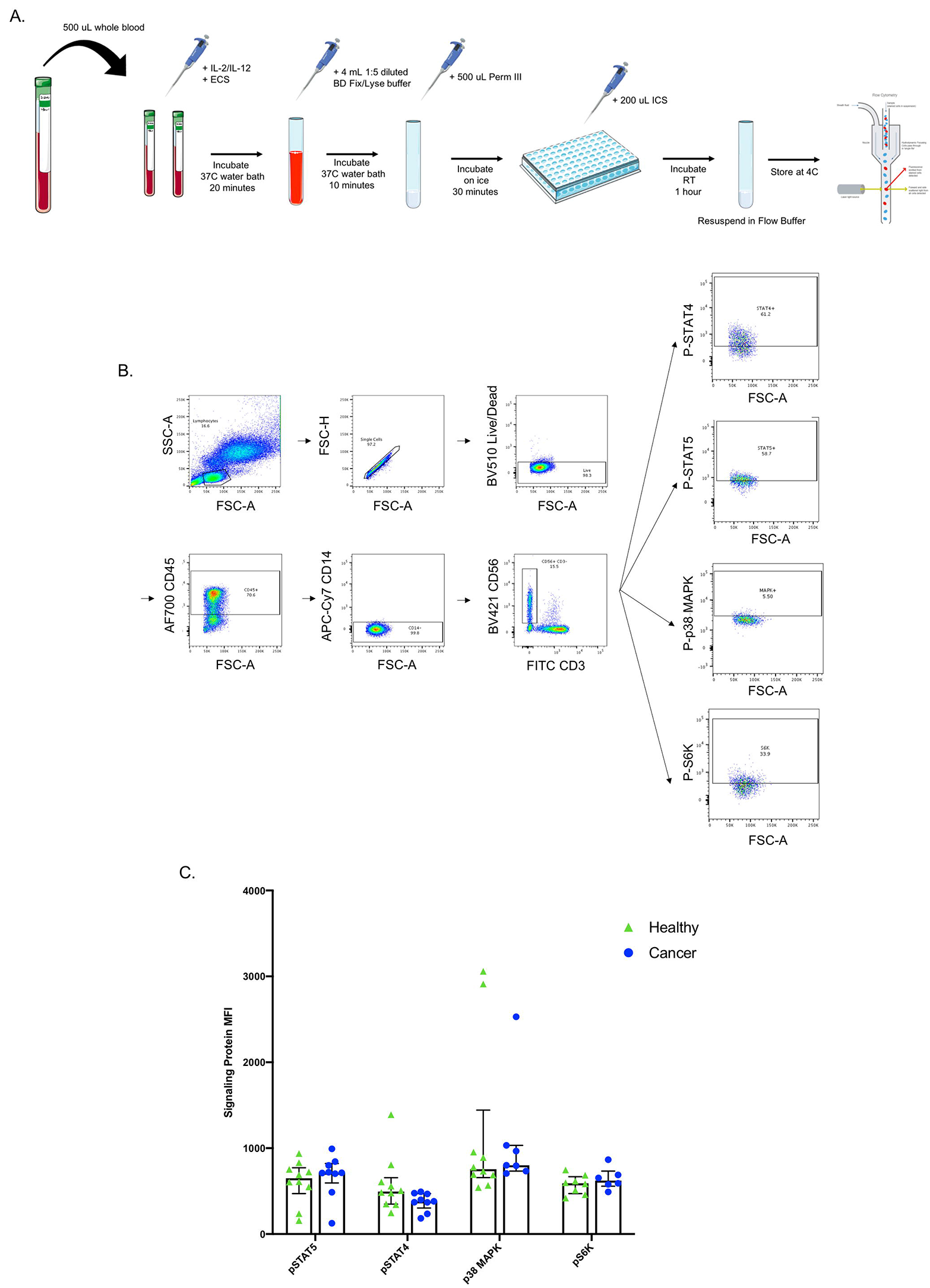
Whole blood signaling protein methodology, gating strategy, and quantification. Whole blood was collected from patients and 200 μL was aliquoted per receptor panel. After a 15-minute incubation with IL−2/12 stimulation and the desired receptor panel, lyse/fix buffer was added and incubated for 10 minutes before blood was spun down. Cells were washed, resuspended in Perm III Buffer, and incubated on ice for 30 minutes before being spun down and resuspended in an intracellular staining mix. Cells were then incubated for an additional hour at room temperature prior to being resuspended and assessed by flow cytometry (**A**). The lymphocyte population was gated on before excluding doublets and dead cells. CD45^+^CD14^−^CD56^+^CD3^−^cells were gated on to assess signaling protein phosphorylation based on unstimulated controls (**B**). The relative level of expression (MFI) of phospho-proteins STAT5, STAT4, p38 MAPK, and S6K was assessed in healthy donor (n=13) and cancer patient (n=9) CD56^+^CD3^−^cells (**C**). Shown are the median values ± IQR.

### 5.3 Protocol 3 – Intracellular IFNγ Staining

#### Quantifying NK cell responsiveness to cytokine stimulation

Finally, NK cell activity was quantified by intracellular and extracellular IFNγ production in response to two stimuli: PMA-ionomycin (a receptor-independent stimulator of cytokine production), and IL−2/12. Intracellular IFNγ was assessed in 11 healthy donors and 9 cancer patients and extracellular IFNγ was assessed in 13 healthy donors and 10 cancer patients. A 6 colour flow cytometry panel and the indicated gating strategy was used to assess the percentage of NK cells producing IFNγ (**Figure 5B**). The healthy donor and cancer patient populations produced intracellular IFNγ in response to both stimuli (**Figure 5C, Supplemental Table 4**). In addition, extracellular IFNγ was quantified in 11 healthy donors and 10 cancer patients from plasma collected prior to intracellular staining. (**Figure 6B, Supplemental Table 4**).

**Figure 5.**
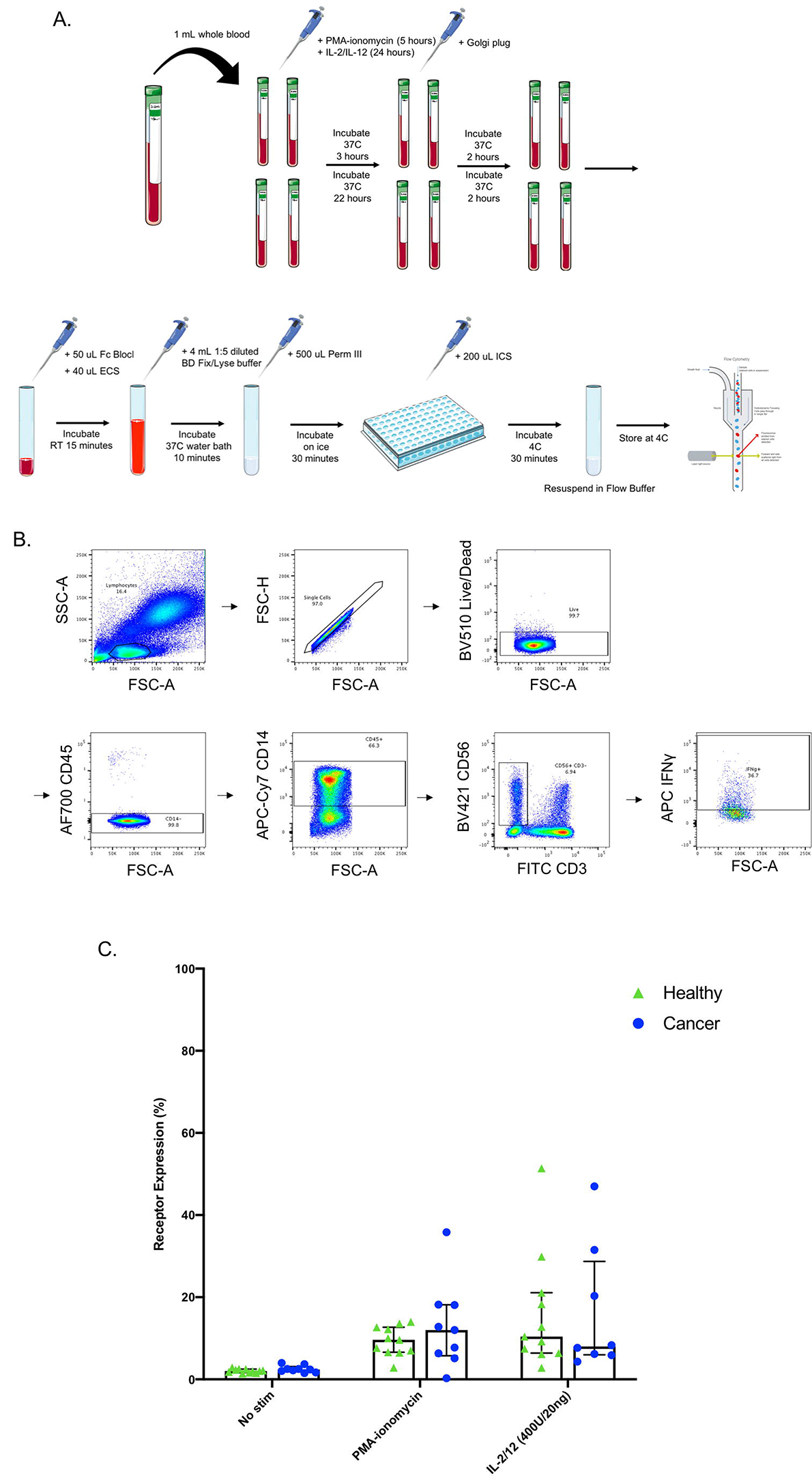
Whole blood secretory cytokine methodology, gating strategy, and quantification. Whole blood was collected from patients and 200 μL was aliquoted per stimulatory condition. Whole blood was stimulated for 24 hours in the presence of PMA-ionomycin, or IL−2/12. At 22 hours post-stimulation, Golgi plug was added to whole blood and incubated for an additional 2 hours. At 24 hours post-stimulation, whole blood was incubated for 15 minutes with an extracellular staining mix. Lyse/fix buffer was then added and incubated for 10 minutes before blood was spun down. Cells were washed, resuspended in Perm III Buffer, and incubated on ice for 30 minutes before being spun down and resuspended in an intracellular staining mix. Cells were then incubated for an additional 30 minutes at 4°C prior to being resuspended and assessed by flow cytometry (**A**). The lymphocyte population was gated on before excluding doublets and dead cells. CD45^+^CD14^−^CD56^+^CD3^−^cells were gated on to assess intracellular IFNγ based on unstimulated controls (**B**). The percentage of CD56^+^CD3^−^cells expressing INFγ after stimulation was assessed in healthy donors (n=11) and cancer patients (n=9) (**C**). Shown are the median values ± IQR.

**Figure 6.**
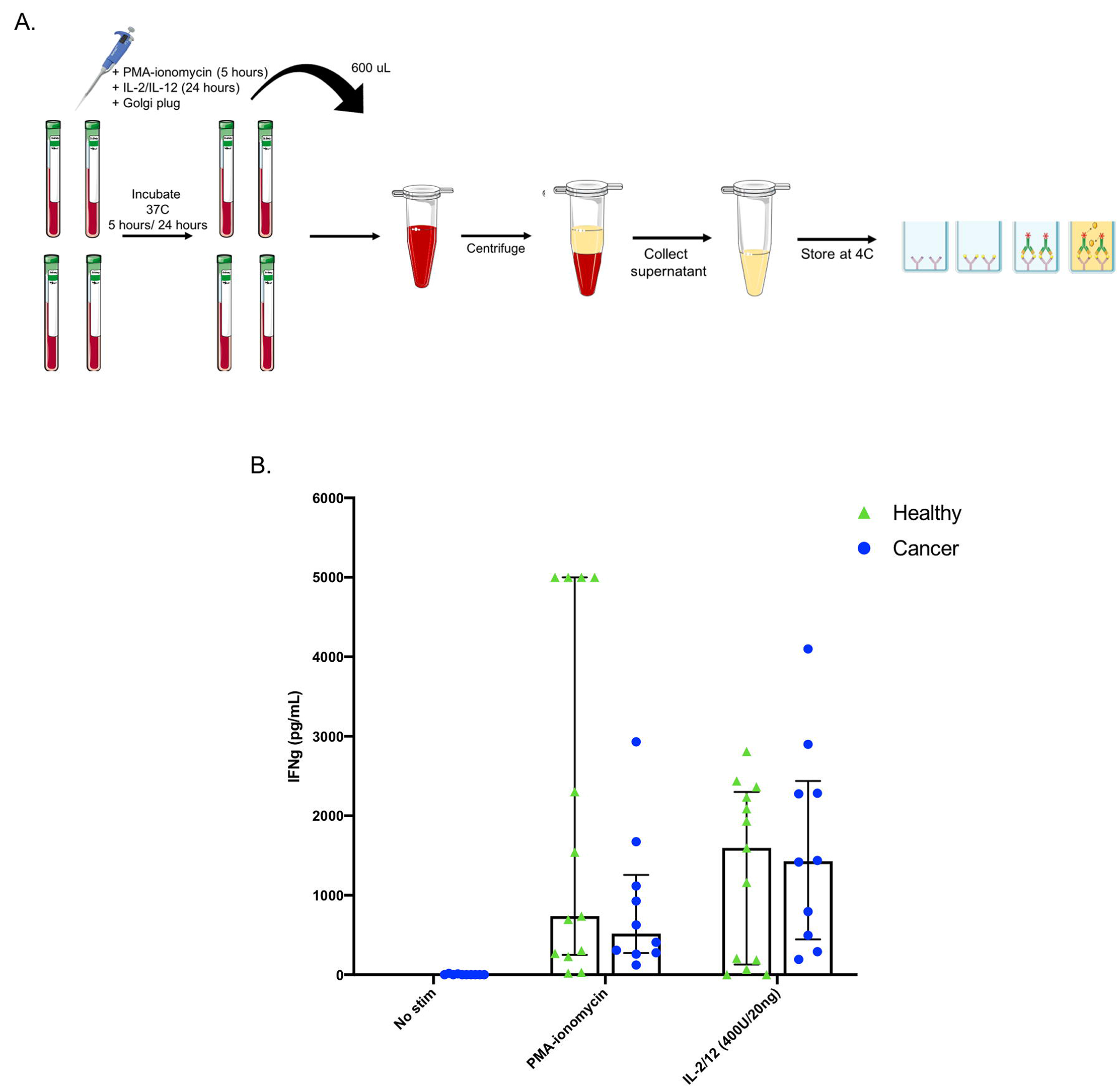
Plasma collection methodology for secretory cytokine quantification by ELISA. After incubation in the absence of stimulation or with either PMA-ionomycin or IL−2/12 stimulation for 24 hours, 600 μL of whole blood was collected and spun down prior to being collected and stored at −80°C for subsequent use in a human IFNγ ELISA (**A**). Extracellular IFNγ was quantified in response to stimuli in healthy donors (n=13) and cancer patients (n=10) (**B**). Shown are the median values ± IQR.

## 6 Advantages

These assays require minimal volumes of blood (200 μL - 1 mL per sample) as compared to methods employing cryopreserved PBMCs (~30−40 mL). This allows for the simultaneous assessment of many targets and the use of whole blood from one patient for multiple assays. In addition, patients may be more likely to consent to blood draws for research that requires minimal blood volumes. These assays can be used to effectively measure extracellular and intracellular targets in healthy as well as disease states (cancer). Here we have assessed NK cell phenotype and function, however, the use of whole blood allows for the assessment of any immune cell, including neutrophils which would otherwise the excluded in PBMC isolation protocols. Finally, a discrepancy can be seen between extracellular targets measured on NK cells using whole blood versus cryopreserved PBMCs. We believe that assessing immune cell phenotypes using whole blood may be more biologically relevant as these protocols minimize the time between blood draw and cell staining and reduce the manipulation of cells that may otherwise impact target expression.

## 7 Limitations

There is potential for inter-assay variability as whole blood samples must be processed immediately, and therefore a large cohort cannot be assessed simultaneously without multiple consenting patients and researchers.

## 8 Troubleshooting

- Antibodies need to be titrated using whole blood to ensure appropriate staining
- May have to adjust incubation times for signaling proteins depending on target (15−30 minutes is a good range to test)
- Mix FACS lyse/fix and blood vigorously to ensure lysis of RBCs; if many RBCs left in pellet, repeat lyse/ fix a second time
- May want to test different Golgi plug timepoints − 4 hours vs. 2 hours
- Intracellular staining can be done in falcon tubes instead of 96 well v-bottom plate, but the pellet is easier to visualize in the plate
- Have also resuspended in PFA instead of flow buffer for longer storage at 4°C.
- CD45 and CD14 staining make the CD56+ CD3-population cleaner but are not necessary
- ELISA: Will need to test different dilutions to prevent OVERFLOW wells − 5x and 10x dilutions are usually acceptable

## 9 Discussion

The workflow of some immune cell studies may be more compatible with protocols utilizing cryopreserved samples, for example multi-institute studies, however, due to the advantages reported here we suggest that some assays may be significantly improved through the implementation of whole blood protocols. These assays circumvent the limitations associated with the use of cryopreserved PBMCs, namely manipulation of cells and the thawing process which may alter cell phenotype and function. Although comparisons have previously been made in the literature between cryopreserved and whole blood PBMC assays, we show that cryopreservation results in an aberrant NK cell phenotype and this is in keeping with what has been previously shown in the literature(24,33,34). We suggest that this discrepancy may lead to misinterpreted conclusions about altered immune cell phenotype and function. As a possible solution, we present methodologies for parallel assessment of immune cell receptor expression, signaling protein activity, and cytokine production in whole blood-derived NK cells. We have demonstrated the feasibility of these assays through the detection of target protein expression in both healthy and disease states (namely solid malignancies), the reproducibility of these assays in patient cohorts despite inherent inter-patient heterogeneity, and the validity of these assays in that our results are comparable to those previously described in the literature(2,12,32,34–39). They are simple, time-efficient, and allow for the assessment of any peripheral immune cell population using a minimal volume of whole blood. Finally, we suggest that they could be used to assess immune cell phenotype and function in any pathological condition, provided sufficient blood volumes.

## 11 Author Contributions

MM and GT performed the experiments and data analysis and were responsible for manuscript preparation. JN and MS screened and consented all patients and performed blood processing on cryopreserved samples. MM, GT, CD, MK, and RA oversaw experimental design and data interpretation.

## 13 Data Availability Statement

The datasets for this study can be made available upon reasonable request.

