## Supplemental Tables for "A Comprehensive Assessment of Human Natural Killer Cell Phenotype and Function in Whole Blood"

### Table 1. Percentage and Median Fluorescence Intensity of NK Cell Surface Receptors

| | % positive CD56+ CD3- cells (median, IQR) | | MFI (median) | |
|---|---|---|---|---|
| **Target** | **Healthy** | **Cancer** | **Healthy** | **Cancer** |
| **NKG2D** | 68.2 (64.1-86.6) | 82.2 (65.4-83.8) | 184.5 (159.5-220.3) | 196.5 (163.5-233.3) |
| **DNAM-1** | 90.4 (88.7-92.5) | 88.4 (83.1-91.6) | 302.0 (258.8-385.3) | 269.5 (233.0-295.0) |
| **NKG2A** | 37.7 (30.3-44.7) | 33.6 (26.9-46.3) | 22.9 (18.2-37.9) | 20.5 (16.7-43.5) |
| **PD-1** | 1.2 (0.6-1.6) | 1.0 (0.6-1.4) | 44.8 (38.7-51.6) | 42.4 (38.9-50.4) |
| **TIGIT** | 33.4 (28.4-44.0) | 33.9 (25.9-42.4) | 271.5 (239.5-288.0) | 262.5 (207.0-292.5) |
| **TIM-3** | 39.6 (29.4-42.3) | 34.8 (28.0-45.9) | 347.5 (303.3-441.8) | 326.5 (255.8-431.3) |
| **CD25** | 7.0 (5.4-9.0) | 7.8 (6.2-10.7) | 82.2 (65.5-177.5) | 110.7 (50.3-195.8) |
| **CD122** | 95.5 (93.7-97.4) | 93.7 (91.5-95.6) | 517.0 (388.5-673.0) | 391.0 (269.5-519.8) |
| **CD132** | 59.0 (47.4-76.3) | 49.1 (40.6-68.2) | 178.0 (136.0-207.0) | 193.0 (181.0-198.0) |
| **CD212** | 36.7 (33.8-38.8) | 43.4 (36.5-51.7) | 200.5 (163.0-215.0) | 200.5 (163.0-215.0) |

### Table 2. Percentage and Median Fluorescence Intensity of NK Cell Activating/ Inhibitory/ Cytokine Receptors Measured in Cryopreserved PBMCs

| | % positive CD56+ CD3- cells (median, IQR) | | MFI (median) | |
|---|---|---|---|---|
| **Target** | **Frozen PBMCs** | **Whole Blood** | **Frozen PBMCs** | **Whole Blood** |
| **NKG2D** | 12.8 (2.6-17.9) | 82.2 (65.4-83.8) | 152.0 (115.5-214.8) | 196.5 (163.5-233.3) |
| **DNAM-1** | 77.5 (64.9-84.1) | 88.4 (83.1-91.6) | 406.5 (310.3-454.8) | 269.5 (223.0-295.0) |
| **NKG2A** | 78.8 (59.6-86.6) | 35.6 (26.9-46.3) | 102.2 (92.0-125.3) | 20.5 (16.7-43.5) |
| **PD-1** | 27.0 (8.7-41.0) | 1.0 (0.6-1.4) | 152.0 (126.8-209.0) | 42.4 (38.9-50.4) |
| **TIGIT** | 94.8 (82.6-97.7) | 33.9 (25.8-42.4) | 320.5 (308.3-409.0) | 262.5 (207.0-292.5) |
| **TIM-3** | 52.8 (44.4-73.9) | 34.8 (28.0-45.9) | 940.5 (846.5-1091.3) | 326.5 (255.8-431.3) |
| **CD25** | 13.1 (11.6-16.8) | 7.8 (6.2-10.7) | 231.0 (221.3-447.5) | 110.7 (50.3-195.8) |
| **CD212** | 32.3 (24.8-39.6) | 43.4 (36.5-51.7) | 519.0 (500.8-554.5) | 200.5 (163.0-215.0) |

### Table 3. Median Fluorescence Intensity of Intracellular Signaling Molecules

| | MFI (median) | |
|---|---|---|
| **Target** | **Healthy** | **Cancer** |
| **STAT5** | 649.5 (563.0-745.3) | 714.0 (703.0-799.0) |
| **STAT4** | 494.5 (381.8-594.0) | 381.0 (37003-467.0) |
| **p38 MAPK** | 755.5 (688.3-938.3) | 801.0 (764.5-999.5) |
| **S6K** | 591.5 (497.0-632.5) | 622.5 (582.0-679.5) |

### Table 4. Intracellular and Extracellular Measurements of IFNγ

| | % positive CD56+ CD3- cells (median, IQR) | |
|---|---|---|
| **Target** | **Healthy** | **Cancer** |
| **Intracellular IFNγ (PMA-ionomycin stimulation)** | 9.61 (6.8-12.5) | 12.0 (6.3-18.1) |
| **Intracellular IFNγ (IL-2/12 stimulation)** | 10.4 (6.9-19.7) | 8.0 (6.1-23.1) |
| | **ELISA (pg/mL)** | |
| **Target** | **Healthy** | **Cancer** |
| **Extracellular IFNγ (PMA-ionomycin stimulation)** | 737.3 (268.2-5000.0) | 516.2 (284.3-1069.7) |
| **Extracellular IFNγ (IL-2/12 stimulation)** | 1596.1 (185.9-2234.3) | 1427.8 (570.9-2281.8) |
